## Supplementary figures and images for "The *Drosophila* hematopoietic niche assembles through collective cell migration controlled by neighbor tissues and Slit-Robo signaling"

### Supplement 1 to Figure 1

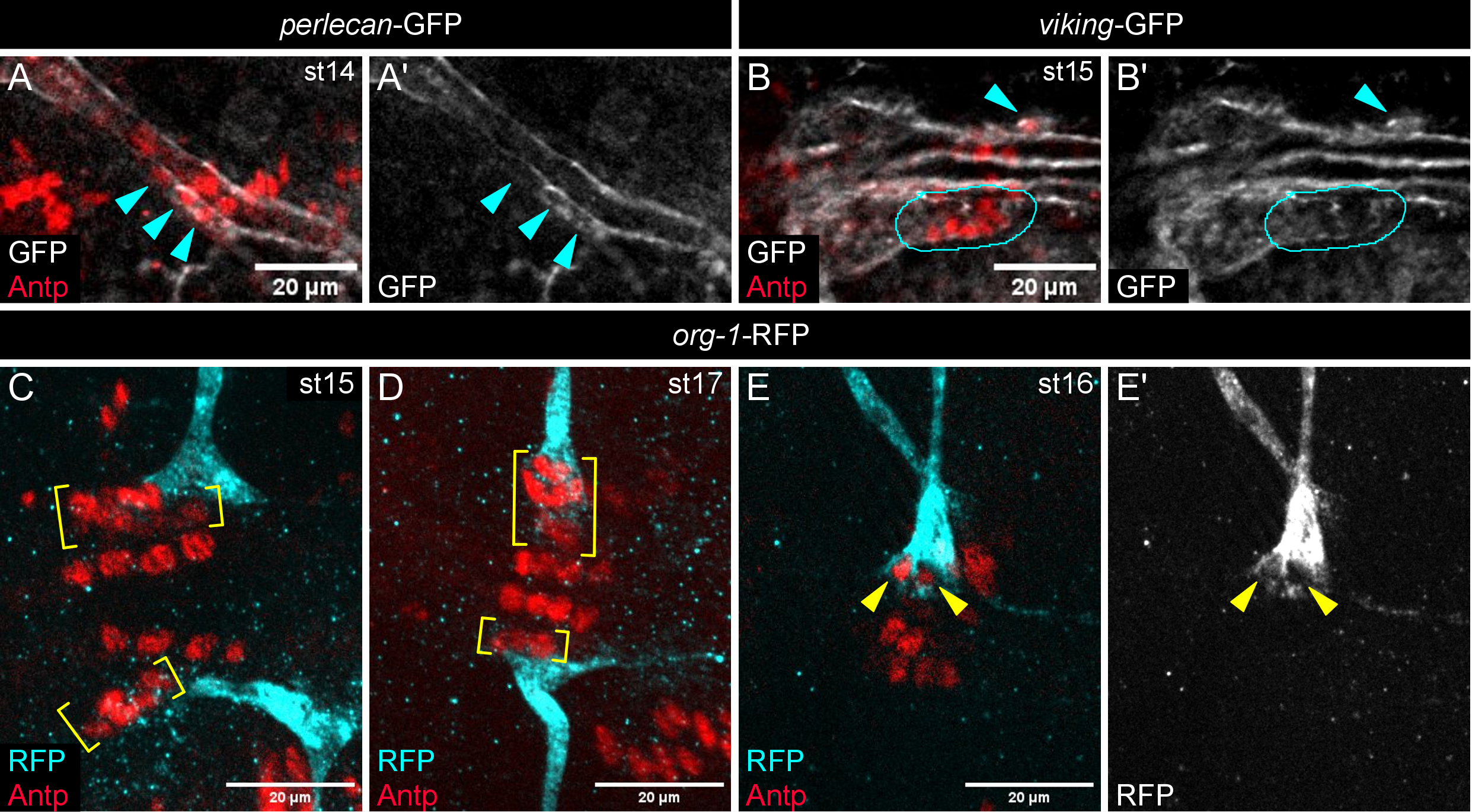

### Supplement 1 to Figure 2

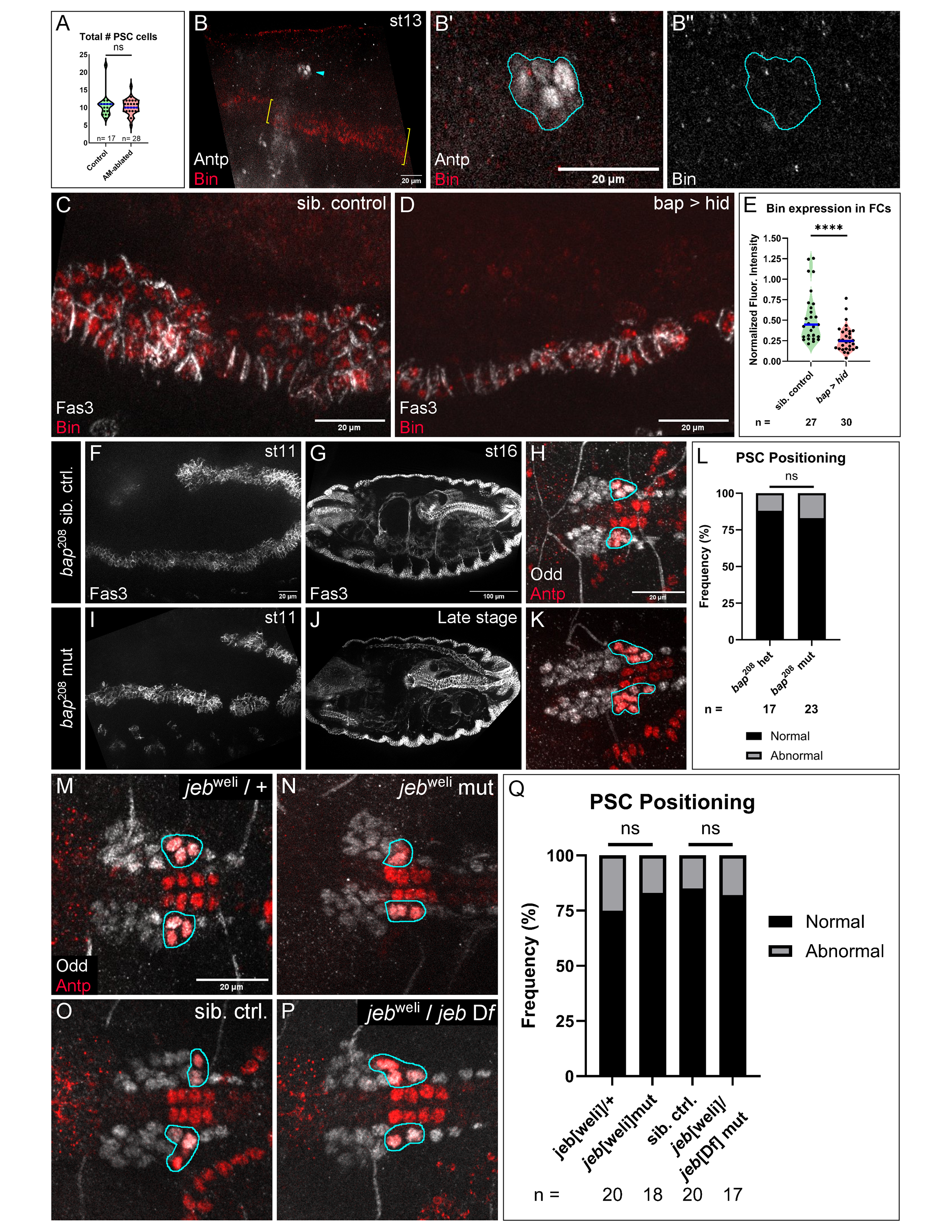

### Supplement 1 to Figure 3

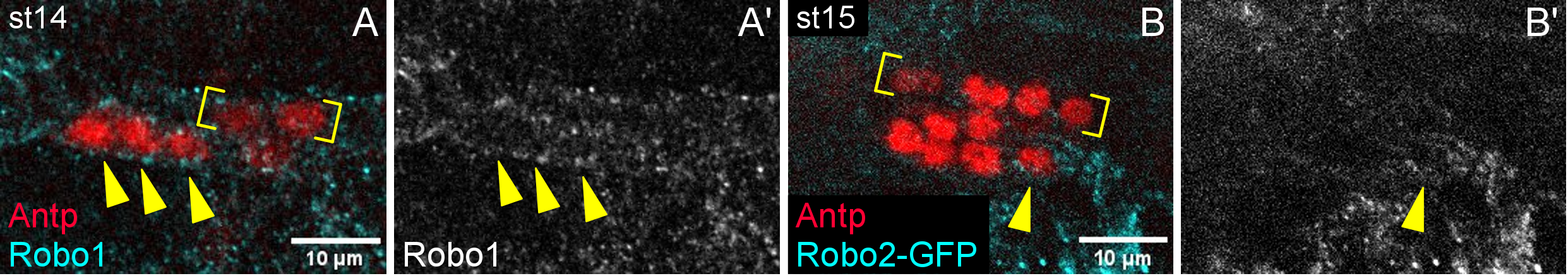

### Supplement 2 to Figure 1

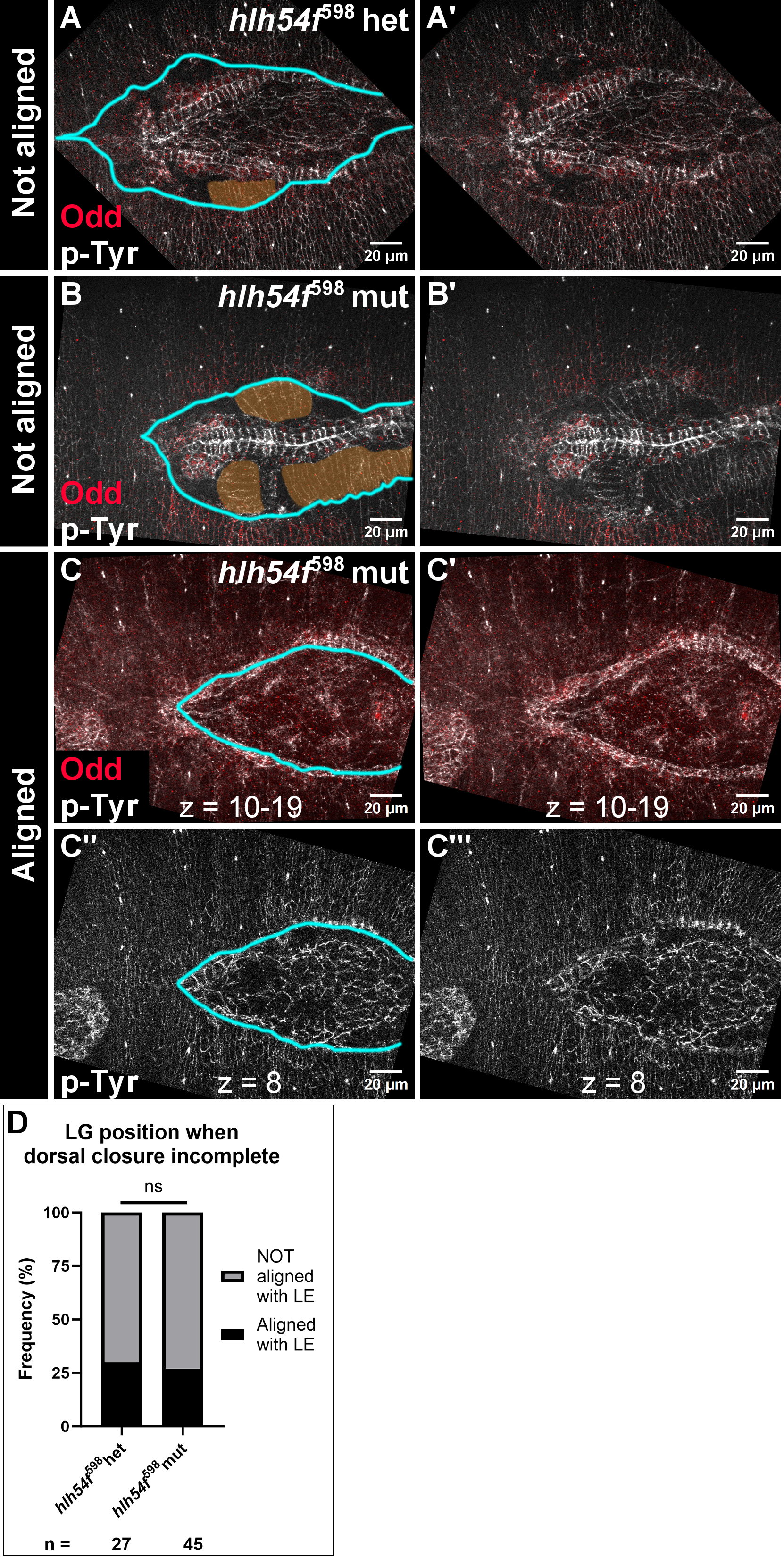
