## Supplementary material for "The *Drosophila* hematopoietic niche assembles through collective cell migration controlled by neighbor tissues and Slit-Robo signaling": Resource Table for Materials

**Key Resources Table**

| **Reagent type (species) or resource** | **Designation** | **Source or reference** | **Identifiers** | **Additional information** |
|---|---|---|---|---|
| genetic reagent (*D. melanogaster*) | Antp-GAL4 | Emerald and Cohen, 2004 | FLYB:FBal0155891 | FlyBase symbol: GAL4$^{Antp\text{-}21}$ |
| genetic reagent (*D. melanogaster*) | *w*$^{1118}$ | Bloomington *Drosophila* Stock Center | BDSC:3605; FLYB:FBal0018186; RRID:BDSC_3605 | FlyBase symbol: w$^{1118}$ |
| genetic reagent (*D. melanogaster*) | Hand-RFP | other | | Gift from Georg Vogler |
| genetic reagent (*D. melanogaster*) | UAS-myr:GFP | Bloomington *Drosophila* Stock Center | BDSC:32200; FLYB:FBti0131976 RRID:BDSC_32200 | FlyBase symbol: P{10XUAS-IVS-myr::GFP}su(Hw)attP1 |
| genetic reagent (*D. melanogaster*) | tupAME-GAL4 | Bataillé et al., 2020 | FLYB: FBtp0142468 | FlyBase symbol:P{tup-GAL4.AME-R}<br>Gift from J.L. Frendo |
| genetic reagent (*D. melanogaster*) | UAS-CD8:GFP | other | | Gift from J.L. Frendo |
| genetic reagent (*D. melanogaster*) | org-1-HN39-RFP | Schaub and Frasch, 2015 | FLYB:FBal0276776 | FlyBase symbol: RFP$^{org\text{-}1.HN39}$ |
| genetic reagent (*D. melanogaster*) | UAS-grim | Hugo Bellen | FLYB:FBti0154788 | Flybase symbol: Dmel\P{UAS-grim.Y}2 |

| | | | | |
|---|---|---|---|---|
| genetic reagent (*D. melanogaster*) | *bin*$^{R22}$ | Zaffran and Frasch, 2001 | FLYB:FBal0043738 | Flybase symbol: Dmel\bin$^{R22}$ |
| genetic reagent (*D. melanogaster*) | *bin*$^{S4}$ | Zaffran and Frasch, 2001 | FLYB:FBal0043739 | Flybase symbol: Dmel\bin$^{S4}$ |
| genetic reagent (*D. melanogaster*) | UAS-hid | Bloomington *Drosophila* Stock Center | BDSC:65403; FLYB:FBti0183136 RRID:BDSC_65403 | Flybase symbol: Dmel\P{UAS-hid.Z}2 |
| genetic reagent (*D. melanogaster*) | bap-GAL4 | Zaffran and Frasch, 2001 | BDSC:91540; FLYB:FBti0214156 | Flybase symbol: Dmel\P{bap-GAL4.3}1.1 |
| genetic reagent (*D. melanogaster*) | bap-GAL4 | other | | gift from Manfred Frasch; Chr: X |
| genetic reagent (*D. melanogaster*) | tinCΔ4-GAL4 | Bloomington *Drosophila* Stock Center | BDSC:92965; FLYB:FBti0216630 | Flybase symbol: Dmel\P{tinC-Gal4.Δ4}12a |
| genetic reagent (*D. melanogaster*) | *slit*$^{2}$ | Bloomington *Drosophila* Stock Center | BDSC:3266 FLYB:FBal0015700 | Flybase symbol:Dmel\sli$^{2}$ |
| genetic reagent (*D. melanogaster*) | *robo1*$^{GA285}$ | other | FLYB:FBal0032588 | Gift from Greg Bashaw Flybase symbol:Dmel\robo1$^{1}$ |
| genetic reagent (*D. melanogaster*) | *robo2*$^{1}$ | Rajagopalan and Dickinson, 2000 | FLYB:FBal0121562 | Gift from Greg Bashaw Flybase symbol:Dmel\robo2$^{1}$ |
| genetic reagent (*D. melanogaster*) | *robo2*$^{123}$ | other | FLYB:FBal0123720 | Gift from Greg Bashaw Flybase |

| | | | | |
|---|---|---|---|---|
| | | | | symbol:Dmel \robo2$^{X123}$ |
| genetic reagent (*D. melanogaster*) | UAS-Slit RNAi #1 | Vienna *Drosophila* Stock Center | VDRC:v108853 FLYB:FBti0159991 | Flybase symbol: Dmel\P{KK100803}VIE-260B |
| genetic reagent (*D. melanogaster*) | UAS-Slit RNAi #2 | Bloomington *Drosophila* Stock Center | BDSC:31468 FLYB:FBal0245521 | Flybase symbol:Dmel \sli$^{JF01229}$ |
| genetic reagent (*D. melanogaster*) | UAS-Robo1 OX | Evans and Bashaw, 2015 | BDSC:97240 FLYB:FBal0316479 | Flybase symbol:Dmel \robo1$^{\Delta C.10xUAS.Tag:HA,Tag:SS(wg)}$ |
| genetic reagent (*D. melanogaster*) | UAS-dcr2 | Bloomington *Drosophila* Stock Center | BDSC:24650 FLYB:FBti0100275 | Flybase symbol:Dmel \P{UAS-Dcr-2.D}2 |
| genetic reagent (*D. melanogaster*) | UAS-dcr2 | Bloomington *Drosophila* Stock Center | BDSC:24651 FLYB:FBti0100276 | Flybase symbol:Dmel \P{UAS-Dcr-2.D}10 |
| genetic reagent (*D. melanogaster*) | svp-lacZ | Bloomington *Drosophila* Stock Center | BDSC:7314 FLYB:FBti0002862 | Flybase symbol: Dmel\P{HZ}svp$^{3}$ |
| genetic reagent (*D. melanogaster*) | perlecan-GFP | Flytrap; GFP Protein Trap Database | FLYB:FBal0243609 | Flybase symbol:Dmel \trol$^{ZCL1700}$ |
| genetic reagent (*D. melanogaster*) | viking-GFP | Buszczak and Spradling, 2007 | FLYB:FBal0211825 | Flybase symbol:Dmel \vkg$^{CC00791}$ |
| genetic reagent (*D. melanogaster*) | *bap*$^{208}$ | Bloomington *Drosophila* Stock Center | BDSC:91539 FLYB:FBal0034201 | Flybase symbol:Dmel \bap$^{208}$ |

| | | | | |
|---|---|---|---|---|
| genetic reagent (*D. melanogaster*) | *jeb*$^{weli}$ | Stute and Holz, 2004 | FLYB:FBal0159133 | Flybase symbol:Dmel\jeb$^{weli}$ |
| genetic reagent (*D. melanogaster*) | *jeb* Df | Bloomington *Drosophila* Stock Center | BDSC:26551 FLYB:FBab0045764 | Flybase symbol:Df(2R)BSC699 |
| genetic reagent (*D. melanogaster*) | robo2-GFP | Bloomington *Drosophila* Stock Center | BDSC:61774 FLYB:FBal0265307 | Flybase symbol:Dmel\robo2$^{MI04295}$ |
| antibody | anti-Antp (Mouse monoclonal) | Developmental Studies Hybridoma Bank | Cat#:8C11, RRID:AB_528083 | IF(1:50) |
| antibody | anti-Odd skipped (Rabbit polyclonal) | Ward and Skeath, 2000 | | IF(1:400); gift from James Skeath |
| antibody | anti-GFP (Chick polyclonal) | Aves labs | Cat#:GFP-1020 RRID:AB_2307313 | IF(1:1500) |
| antibody | anti-Fas3 (Mouse monoclonal) | Developmental Studies Hybridoma Bank | Cat#:7G10 RRID:AB_528238 | IF(1:50) |
| antibody | anti-Mef2 (Rabbit polyclonal) | Developmental Studies Hybridoma Bank | Cat#:Mef2 RRID:AB_2892602 | IF(1:1000) |
| antibody | anti-Slit (Mouse monoclonal) | Developmental Studies Hybridoma Bank | Cat#:C555.6D RRID:AB_528470 | IF(1:200); gift from Greg Bashaw |

| | | | | |
|---|---|---|---|---|
| antibody | anti-LacZ (Chick polyclonal) | Abcam | Cat#:ab9361 RRID:AB_307210 | IF(1:1000) |
| antibody | anti-RFP (Rabbit polyclonal) | Abcam | Cat#:ab62341 RRID:AB_945213 | IF(1:1000) |
| antibody | anti-Bin (Rabbit polyclonal) | other | | IF(1:100); gift from Eileen Furlong |
| antibody | anti-Robo1 (Mouse monoclonal) | other | | IF(1:200); gift from Greg Bashaw |
| antibody | anti-Odd skipped (Guinea pig polyclonal) | other | | IF(1:1200); gift from John Reinitz |
| chemical compound, drug | Paraformaldehyde | Electron Microscopy Sciences | Cat#:15710 | |
| chemical compound, drug | Propyl-gallate | Sigma Aldrich | PubChem Substance ID:24898394; SKU:P3130; CAS Number:121-79-9 | |
| chemical compound, drug | Normal Donkey Serum | Jackson ImmunoResearch Labs Inc | Cat#:017-000-121 RRID:AB_2337258 | |
| chemical compound, drug | Ringer’s solution | other | | Recipe from https://doi.org/10.1242/dev.125.15.2781 |

| | | | | |
|---|---|---|---|---|
| chemical compound, drug | Triton X-100 | MilliporeSigma | CAS Number: 9036-19-5 | |
| software, algorithm | FIJI | ImajeJ | RRID:SCR_002285 | http://fiji.sc |
| software, algorithm | Photoshop | Adobe | RRID:SCR_014199 | https://www.adobe.com/products/photoshop.html |
| software, algorithm | Prism | Graphpad | RRID:SCR_002798 | v9.0.0-v10.0.0 |
| software, algorithm | Axio-Vision Imaging Software | Zeiss | | v4.8.1 |
| software, algorithm | VisiView | Visitron | | |
| software, algorithm | Metamorph Microscopy Automation and Image Analysis Software | Leica | | v7.8.40 |
| other | 63x / 1.2 NA water immersion objective | Leica | | |
| other | 60x / 1.3 NA silicone immersion objective | Olympus | | |
| other | AxioCam HRm | Zeiss | | |
| other | 40x / 1.2 NA water immersion objective | Zeiss | | |
| other | 20x / 0.8 NA objective | Zeiss | | |

| | | | |
|---|---|---|---|
| other | M165FC | Leica | |
| other | Achromat 1.6x objective | Leica | |
| other | GFP Filter set ET470/40x; ET525/50m | Leica | |
| other | mCherry Filter set ET560/40x; ET630/75m | Leica | |
| other | pco.edge 4.2 bi sCMOS | PCO | |
| other | Cell Center Stockroom (Penn) | other | RRID:SCR_022399 |
| other | CDB Microscopy Core (Penn) | other | RRID:SCR_022373 |
